# The mouse mammary tumor virus intasome exhibits distinct dynamics on target DNA

**DOI:** 10.1101/2021.11.17.468995

**Authors:** Laura E. Baltierra-Jasso, Nathan D. Jones, Allison Ballandras-Colas, Alan N. Engelman, Richard Fishel, Kristine E. Yoder

**Affiliations:** Department of Cancer Biology and Genetics, The Ohio State University College of Medicine, Columbus, Ohio, USA; Comprehensive Cancer Center, The Ohio State University; Center for Retrovirus Research, The Ohio State University; Institut de Biologie Structurale, Grenoble, France; Department of Cancer Immunology and Virology, Dana-Farber Cancer Institute, Boston, Massachusetts, USA; Department of Medicine, Harvard Medical School, Boston, Massachusetts, USA

## Abstract

Retroviral intasomes are complexes assembled from purified integrase (IN) and oligonucleotides mimicking viral DNA ends (vDNA). Recombinant intasomes faithfully recapitulate integration of vDNA into a target DNA. Structural studies of retroviral intasomes have revealed an array of IN oligomer forms, which appear to share a conserved intasome core coordinating the vDNA ends for strand transfer into the target DNA. Here we have explored the biochemical and dynamic properties of the mouse mammary tumor virus (MMTV) octameric intasome. We show that the MMTV intasome is remarkably stable compared to the prototype foamy virus (PFV) tetrameric intasome. MMTV integration activity peaks within the range of physiological ionic strength and is more active in the presence of manganese compared to magnesium. Single-molecule images demonstrate that the target DNA search by MMTV intasomes appears rate-limiting, similar to PFV intasomes. The time between strand transfer of the two MMTV vDNA ends into the target DNA is ∼3 fold slower than PFV intasomes. MMTV intasomes can form extremely stable, largely immobile filaments on a target DNA that are comprised of multiple intasomes. This unusual property suggests that MMTV intasomes may readily form higher order oligomers that might underpin their increased stability.

## INTRODUCTION

Retroviral RNA genomes are copied to linear double stranded DNA (cDNA) by reverse transcription (1). Integration of this cDNA into host chromatin is an essential step in the retroviral life cycle and is catalyzed by the viral protein integrase (IN) (1-3). During infection retroviral IN performs two catalytic activities. First, a 3’ processing activity cleaves two terminal nucleotides from the cDNA 3’ ends generating terminal recessed hydroxyl groups (3). Second, a strand transfer activity covalently joins the recessed 3’ hydroxyls to host DNA (3-6). Depending on the retrovirus, the two strand transfer events are separated by 4-6 bp of host target DNA. The result is the integrated proviral genome flanked by 4-6 nt single strand gaps of host DNA and 5’ dinucleotide flaps of viral cDNA. Host enzymes appear to repair this gapped integration intermediate resulting in a stable integrated provirus flanked by duplications of host DNA (7).

Retroviral INs have three common domains: an amino terminal domain (NTD), catalytic core domain (CCD), and carboxyl terminal domain (CTD) (reviewed in (3)). Some retroviruses, including the spumavirus prototype foamy virus (PFV), have an additional amino terminal extension domain (NED). The CCD includes a DDE catalytic triad that coordinates divalent metal cations. All domains participate in multimerization to form an integration complex. The linker region between the IN CCD and CTD varies in length between retroviruses and is predictive of the number of INs necessary for an active complex (8-12).

A multimer of retroviral IN binds the viral cDNA ends to form a functional enzymatic complex. This intasome may be assembled *in vitro* with recombinant IN and oligomers mimicking the viral cDNA ends (vDNA) (13). In addition, the vDNA may be labeled with small fluorophores or biotin moieties without affecting integration activity (14). The first retroviral intasome structures revealed a tetramer of PFV IN alone or bound to a target DNA (13,15). The intasome-bound target DNA was distorted to accommodate strand transfer at 4 bp spacing across the major groove of host DNA (15). Two “inner” PFV IN subunits are catalytically active and bind both the vDNA and target DNA. Coordinates for only the CCD of the two “outer” PFV IN protomers have been resolved, which appear important for intasome structure but not catalysis (15-18). Function(s) for the unresolved domains of the outer PFV IN protomers are unknown.

Intasome structures of orthoretroviruses showed a diversity of IN oligomerization (8,9,11-13,15,17,19). The deltaretrovirus human T cell leukemia virus (HTLV-1) intasome is also a tetramer of IN (20,21). The alpharetrovirus Rous sarcoma virus (RSV) and the betaretrovirus mouse mammary tumor virus (MMTV) intasomes are octamers of IN (8,9,12). The lentivirus Maedi visna virus (MVV) intasome is a hexadecamer of IN (8,9,12). Tetramer and dodecamer HIV-1 IN architectures, as well as filaments of intasomes, were observed by single particle cryogenic electron microscopy (11,22,23). All of these intasome structures notably display a conserved intasome core (CIC) that is similar to the central structural elements of the PFV IN tetramer (24).

Retroviral integration in cells is inefficient, making interrogation of integration processes *in vivo* challenging (25-27). Real-time single-molecule imaging has been utilized to dissect the search and integration dynamics of PFV intasomes with target DNA *in vitro* (18,28). These studies demonstrated a target DNA search for 2.1 s while in continuous contact with the DNA backbone, suggesting that PFV intasomes could interrogate ∼1.6 kb of duplex DNA, most likely by rotation coupled diffusion. Once an integration site was detected, the time between the two vDNA strand transfer events was 0.47 s (18,28,29). Remarkably, PFV intasomes performed 100-300 search events per strand transfer event strongly suggesting that target site selection was rate-limiting (28). To date, the dynamic interactions with target DNA have only been described for PFV intasomes (18,28). Whether the CIC mediates similar kinetics with different oligomeric forms of IN protomers, similar to intasomes derived from highly pathogenic betaretroviruses and lentiviruses, is unclear.

While some retroviruses employ a host protein as an integration co-factor, none has been reported for MMTV (30-34). Retroviral integration host co-factors help determine the genomic integration profile relative to chromatin elements such as transcription units, promoters, or CpG islands (35-37). MMTV displays minimal preference for genomic elements, which agrees with the absence of an integration co-factor (30,38,39). Despite this integration profile, the MMTV provirus is able to dysregulate cellular proto-oncogene expression leading to mammary adenocarcinomas in mice. Thus, MMTV infection has been employed as a model for breast cancer pathogenesis (31-33). Integration host co-factors are known to aid in the stability of recombinant intasomes, as in the case of the LEDGF/p75 host co-factor for lentiviruses MVV and HIV-1. However, PFV and MMTV intasomes may be readily assembled from recombinant IN and vDNA only (8,9,11,15,40).

Here we have examined the biochemical properties, strand transfer kinetics, and search dynamics of the MMTV intasome. We found that these octameric intasomes display a longer time between the two strand transfer events and are capable of a prolonged search on target DNA compared to PFV intasomes. When multiple MMTV intasomes are present they appear to form filaments on the target DNA suggesting higher order interactions. These observations are consistent with the conclusion that retroviral intasomes display a common search mechanism with diverse search parameters.

## MATERIALS AND METHODS

### Chemicals

Chemicals were at least 98% pure and were purchased from Sigma-Aldrich, Millipore-Sigma, GoldBio, VWR and ThermoFisher. NHS ester Cy3 and Cy5 dyes were purchased from Lumiprobe. DNA oligonucleotides were synthesized by IDT (Newark, NJ).

### MMTV integrase purification and labeling of viral DNA

The MMTV IN expression construct included an N-terminal hexa-histidine tag, thrombin site, and human rhinovirus (HRV) 3C protease site. The protein was induced with 0.25 mM IPTG in *E*.*coli* Rosetta BL21(DE3) (Novagen) in the presence of 50 µM ZnCl_2_ at 37° C overnight. Bacteria were lysed in 20 mM HEPES (pH 7.5), 1 M NaCl, 5 mM CHAPS and 1 mM PMSF. Cells were sonicated and centrifuged at 120,000 x g for 1 h at 4° C. The supernatant was applied to Ni-NTA Superflow resin (Qiagen) and proteins eluted with a gradient of 20-200 mM imidazole (pH 8.0). The histidine tag was removed by cleavage with HRV 3C protease overnight at 4° C. MMTV IN was further purified with heparin sepharose (GE Healthcare). Pooled fractions were dialyzed in 20 mM HEPES (pH 7.5), 1 M NaCl, 5 mM CHAPS, 2 mM DTT, 0.5 mM EDTA and 10 % glycerol. Catalytically inactive MMTV IN(D122N) was prepared by the same purification method. Oligonucleotides of the transferred strand 5’ CAGGT*CGGCCGACTGCGGCA 3’ and non-transferred strand 5’ AATGCCGCAGTCGGCCGACCTG 3’ mimic the MMTV U5 end. T* indicates the position of a 4-amino-thymine used for conjugation of a Cy3 or Cy5 NHS ester. Labeled oligonucleotides were purified by HPLC Poroshell 120 EC-C18 reverse phase column (Agilent) and 12% urea PAGE. The purified oligonucleotide was annealed to the non-transferred oligonucleotide yielding a preprocessed vDNA.

### MMTV intasome assembly

Intasomes were assembled as previously described (8). Briefly, a 3:1 ratio of purified MMTV IN:vDNA in 20 mM HEPES (pH 7.5), 600 mM NaCl, 2 mM DTT was dialyzed overnight at 4° C against 25 mM Tris-HCl (pH 7.5), 80 mM NaCl, 2 mM DTT, 25 µM ZnCl_2_, 10 mM CaCl_2_. The NaCl concentration was increased to 250 mM and incubated 1 h on ice. Intasomes were purified by size exclusion chromatography with a Superdex 200 Increase (10/300) column (GE Healthcare) equilibrated in 25 mM Tris-HCl (pH 7.5), 200 mM NaCl, 2 mM DTT, 25 µM ZnCl_2_, 10 mM CaCl_2_ and 10% glycerol. Individual fractions were frozen with liquid nitrogen and stored at −80° C. Frozen MMTV intasomes retained activity for at least 6 months.

### Integration assays

Integration reactions were 20 mM HEPES (pH 7.5), 110 mM NaCl, 4 µM ZnCl_2_, 10 mM DTT, 20 mM MgSO_4_, 1.8 nM 3 kb supercoiled pGEMT plasmid (Promega), and 20 nM MMTV intasomes in a final volume of 15 µL. Excluding kinetic assays, all integration reactions were incubated for 1 h at 37° C. Reactions were stopped with the addition of 0.5 % SDS, 0.5 mg/mL proteinase K, 25 mM EDTA (pH 8.0) and incubated for 1 h at 37° C. 18 fmol of a 717 bp linear DNA was added to some reactions as a gel loading control. Integration products were resolved by 1.25 % agarose gel electrophoresis, stained with ethidium bromide and scanned for Cy5 and ethidium bromide fluorescence (Sapphire Biomolecular Imager, Azure Biosystems). Images were quantified with GelAnalyzer software (Azure Biosystems). Cy5 fluorescent products include unreacted vDNA, concerted integration (CI^1^) of both vDNA resulting in linear products, and half site integration (HSI) of a single vDNA to the target DNA yielding a relaxed circle. Ethidium bromide images visualize the unreacted supercoiled target DNA, linear and relaxed plasmid DNA, and the loading control. Multiple CI events to a single supercoiled target DNA resulted in a Cy5 smear (Cl^2^ products). The Cl^2^ products were included in the calculation of total CI (Cl = CI^1^ + Cl^2^) as previously described (29). All experiments were performed in triplicate with at least two independent preparations of MMTV intasomes. Statistical significance of integration reactions was determined by paired t test to generate two-tail P values (Excel).

### Aggregation assays

20 nM MMTV intasomes were incubated in 20 mM HEPES (pH 7.5), 110 mM NaCl, 4 µM ZnCl_2_, 10 mM DTT, 20 mM MgSO_4_, except where noted, for 1 h at 37° C in 100 µL total volume. Where indicated, 1.8 nM linear 3 kb DNA was included. Samples were centrifuged at 18,000 x g for 30 min at 4° C. Pellets were resuspended in denaturing PAGE buffer and resolved by 12% PAGE. Gels were stained with Coomassie Brilliant Blue, scanned (Sapphire Biomolecular Imager, Azure Biosystems), and quantified (GelAnalyzer software, Azure Biosystems). Band intensities were normalized to the sample without MgSO_4_ or the sample with the lowest concentration of NaCl. Statistical significance of aggregation reactions was determined by paired t test to generate two-tail P values (Microsoft Excel).

### Single-molecule magnetic tweezers

Single-molecule magnetic tweezers analysis was performed as previously described (28,29). Briefly, a 7 kb DNA substrate was generated by ligating a linker with multiple biotins to one end and a linker with multiple digoxygenins to the other end. Glass slides were treated with 3-aminopropyl triethoxysilane and passivated with a biotin-PEG SVA/mPEG SVA mix (Invitrogen). Treated glass slides were assembled with double sided tape and an aluminum backing to generate flow channels in a custom microscope slide. The DNA substrate was tethered to the surface via NeutrAvidin (Invitrogen). Paramagnetic beads (Thermo Fisher Scientific) coated with anti-digoxygenin (Novus Biologicals) were injected into the channel. Ten negative supercoils were induced by an equivalent number of counter-clockwise turns of a neodymium magnet above the flow cell slide at 0.3 pN force. Reactions containing 20 nM MMTV intasomes in 30 mM Bis-tris propane (pH 7.5), 110 mM NaCl, 4 µM ZnCl_2_, 250 µM DTT, 20 mM MgSO_4_, 200 μg/mL acetylated BSA, 0.02 % IGEPAL were flowed into the channel. Movies were recorded for 30 min at a 100 ms frame rate. The 3D positions of the paramagnetic beads were evaluated using the tracking software Video Spot Tracker (CISMM at UNC-CH). Resultant coordinates were analyzed using custom MATLAB scripts (MathWorks). The time between the two strand transfer events (*τ*_ST_) was determined for concerted integration events; the initial change in the z-position corresponded to the first strand transfer and subsequent movement in all axes (x-, y- and z-) indicated the second strand transfer. Histograms generated from n events were fit as a single exponential decay to determine the mean *τ*_ST_ and standard error (Origin, OriginLabs). Binning of histograms was performed as described previously with a bin minimum of 100 ms (28,41,42).

### Single-molecule total internal reflection fluorescence

Single-molecule total internal reflection fluorescence (smTIRF) was performed as previously described (28). Briefly, phage λ DNA was digested with restriction endonuclease XmaJI to generate 24 kb fragment DNAs. Biotin labeled DNA linkers were ligated to the fragment ends. Flow cells were assembled from quartz slides treated with 1 biotin-PEG-SVA:300 mPEG-SVA and glass cover slips treated with mPEG-SVA. NeutrAvidin was injected into the flow cell followed by addition of biotinylated λ fragments stretched by flow.

Imaging was performed with 50-100 pM or 2 nM MMTV intasomes with Cy5 or Cy3 labeled vDNA in 30 mM Bis-tris propane (pH 7.5), 110 mM NaCl, 20 mM MgSO_4_, 4 µM ZnCl_2_, 100 µM DTT, 200 μg/mL acetylated BSA, 0.02 % IGEPAL, 20 nM protocatechuate 3,4-dioxygenase (PCD), and 5 mM protocatechuic acid. PCD was prepared as described (43,44). Fluorescence was detected with a custom built prism TIRF microscope (Olympus IX-71, water-type 60X objective NA = 1.2, 1.6X extended magnification) and recorded on an electron-multiplying charge-coupled device camera (EMCCD, Princeton Instruments, ProEM 512 excelon). Lifetime and diffusion were recorded for 1200 s at a 250 ms frame rate. Following the real-time recordings, the target DNA was stained with Syto 59 Red Fluorescent Nucleic Acid Stain (Thermo Fisher Scientific) and the images overlaid onto the Cy5 or Cy3 movies. Particles were tracked using DiaTrack software (Sydney, Australia) and the results were analyzed with MATLAB (MathWorks) and Origin Pro (OriginLab) (45).

## RESULTS

### Purified MMTV intasomes catalyze concerted integration

Recombinant MMTV intasomes were assembled with Cy5 labeled vDNA (8,9,13,15,17). Size exclusion chromatography (SEC) revealed a high molecular weight peak (530 kDa) consistent with an asymmetric octamer of MMTV IN and two vDNAs (Supplementary Figure S1A). Two additional peaks were also observed, consistent with a dimer of MMTV IN (100 kDa) and free vDNA (19 kDa).

MMTV intasome SEC fractions were analyzed for integration activity by agarose gel electrophoresis. Concerted integration (CI^1^) of the two vDNAs to a supercoiled (SC) plasmid results in a linear product (LN) (Supplementary Figure S1B). Additional concerted integration (CI) into the linear CI^1^ product may produce an array of shortened linear products (Cl^2^). Intasomes may also integrate a single vDNA to the target plasmid, termed half site integration (HSI), appearing as a product with the mobility of a relaxed circle (RC). Gels were visualized by ethidium bromide staining of DNA and Cy5 fluorescence of vDNA and integration products (Supplementary Figure S1C, D). Analysis of SEC fractions indicated that the octameric MMTV intasome was active, which can be easily observed by the generation of Cy5 fluorescent CI^1^ and Cl^2^ products.

Divalent cations are required for the assembly and catalytic activities of retroviral integration complexes (46-48). HIV-1 IN is catalytically active in the presence of magnesium or manganese, but is not active in the presence of calcium (46,48,49). This observation provided the foundation for the purification of intasomes in the presence of calcium, effectively preventing catalysis until the addition of magnesium or manganese (8,50).

The effects of magnesium and manganese on MMTV intasome integration activity were examined in reactions that included 1 mM CaCl_2_ which was carried over from the SEC purification buffer. Cy5 labeled CI^1^ products were observed beginning at a 2 fold molar excess of MgCl_2_ to CaCl_2_, with CI^1^ products continuing to increase up to the maximum of 15 fold molar excess used in these experiments (Figure 1A, Supplementary Figure S2A). At 10 mM MgCl_2_ we observed the accumulation of Cl^2^ products. Autointegration (AI) products are the result of one vDNA integrating to a second vDNA. AI products were observed in the presence of 15 mM MgCl_2_. In the presence of 1 mM MnCl_2_ we observed Cl^1^, Cl^2^ and HSI products. (Figure 1B, Supplementary Figure S2B). The increased integration activity in the presence of MnCl_2_ likely reflects an enhanced ability of manganese ions to displace calcium ions, similar to previous studies with isocitrate dehydrogenase and chelators such as EGTA (51). However, MnCl_2_ also substantially increased the fraction of AI products, which could potentially complicate biochemical and kinetic analysis. In addition, we have found that manganese is incompatible with the oxygen scavenging system (OSS) that is essential for single-molecule imaging analysis. Equivalent concentrations of MgSO_4_ yielded approximately 30% less CI products compared to MgCl_2_ (Figure 1C, Supplementary Figure S2C). However, MgSO_4_ resulted in virtually undetectable AI products and was used in subsequent reactions. The biochemical cause(s) of the chloride versus sulfate anion dissimilarity in MMTV integration is unknown.

**Figure 1.**
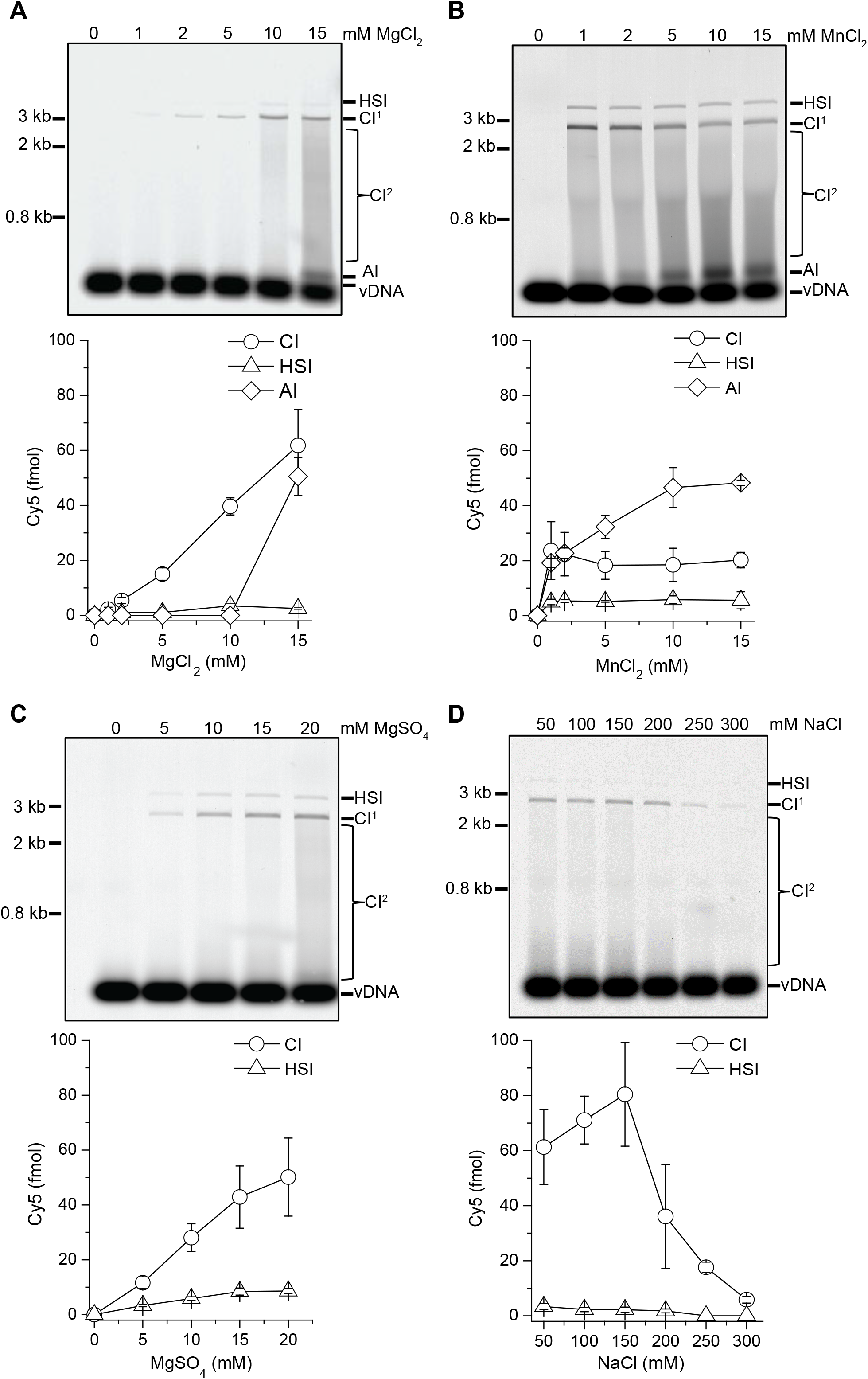
MMTV intasomes have different integration efficiencies in the presence of manganese or magnesium. MMTV intasome integration to a SC plasmid was performed with increasing concentrations of (**A**) MgCl_2_, (**B**) MnCl_2_, (**C**) MgSO_4_, and (**D**) NaCl. Integration products were separated by agarose gel electrophoresis and quantified by the Cy5 fluorescent signal in each lane (fmol). Half-site integration (HSI) products where only one vDNA has been joined to the plasmid have the mobility of a relaxed circle. A single concerted integration product (CI^1^) has the mobility of a linear plasmid. Additional CI events (CI^2^) result in a smear between CI^1^ and unreacted vDNA. Autointegration (AI) of vDNA to vDNA has slower mobility than unreacted vDNA. All reactions include 1 mM CaCl_2_ which is present during intasome assembly and purification. MMTV intasomes are not active in the absence of magnesium or manganese divalent ions. Increasing concentrations of MgCl_2_ or MgSO_4_ lead to increasing total CI (CI = CI^1^ + CI^2^). AI products were only apparent at the highest concentration of MgCl_2_. In contrast, MMTV intasome CI was equivalent at all concentrations of MnCl_2_ assayed, while AI products increased with increasing concentrations of MnCl_2_. HSI products were unaffected by the increasing concentrations of divalent ions. MMTV intasomes were also assayed with a titration of NaCl in the presence of 20 mM MgSO_4_. MMTV CI products peaked in the physiologically relevant range of 100-150 mM NaCl. Fewer CI products were observed at >150 mM NaCl. MMTV integration efficiency appeared to be reduced at 50 mM NaCl, a concentration less than physiological, but was not statistically significant. Error bars indicate the standard deviation of at least 3 independent experiments performed with at least 2 independent MMTV intasome preparations.

### MMTV intasomes function best at physiologically relevant ionic strength

The integration activity of MMTV intasomes was assessed over a range of NaCl concentrations (Figure 1D, Supplementary Figure S2D). The formation of CI products peaked at 150 mM NaCl and appeared to rapidly decrease to nearly undetectable at 300 mM NaCl. We conclude that that the maximal MMTV activity occurs within the physiological range of ionic strength. All subsequent biochemical, kinetic and single-molecule studies of MMTV intasome activity were performed in 110 mM NaCl, in a final reaction buffer composition similar to physiological ionic strength.

### MMTV intasome integration displays saturation kinetics

MMTV intasome integration to a SC plasmid target DNA was measured over time. MMTV intasomes generated CI products for at least 80 min with product saturation occurring when only 50-60% of the target SC plasmid remained (Figure 2A; Supplementary Figure S2E). In contrast, HSI products reached a plateau after 5 min and remained at ∼10% of the total integration products. These results appear consistent with the conclusion that HSI mostly results from a fraction of defective intasomes rather than a kinetically separable intermediate that progresses to a CI product.

**Figure 2.**
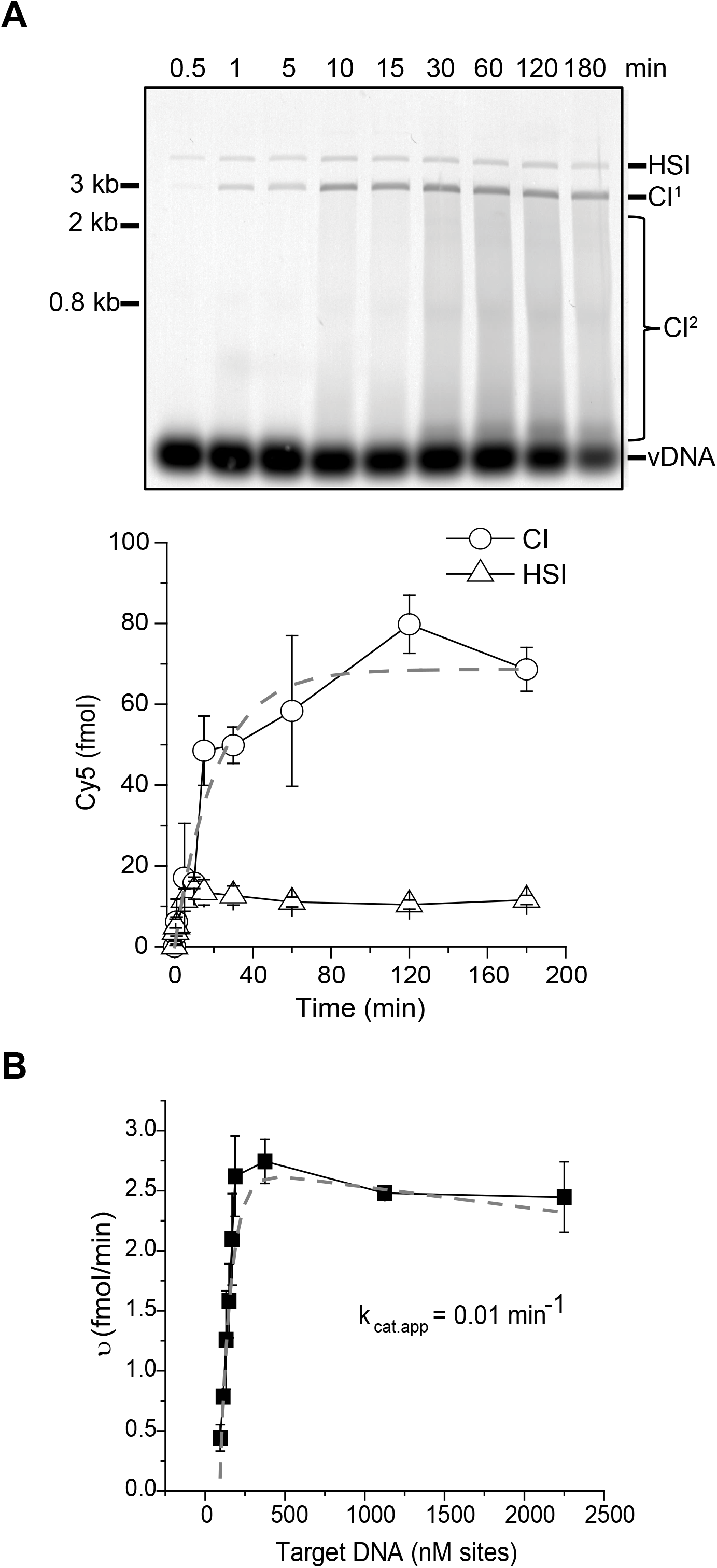
Time course and catalytic rate of MMTV intasomes. (**A**) MMTV intasomes were evaluated for integration to a SC plasmid for up to 3 h. Integration products were separated by agarose gel electrophoresis and quantified by the Cy5 fluorescent signal in each lane (fmol). Based on the accumulation of Cy5 CI products, MMTV intasomes retained activity for at least 80 min. (**B**) The initial rate of reaction was estimated based on velocity vs substrate concentration. Increasing SC plasmid concentrations were converted to nM sites (0.25, 0.3, 0.35, 0.4, 0.45, 0.5, 1, 3 and 6 nM) considering the number of possible MMTV intasome binding sites in the 3 kb target plasmid. The MMTV intasome minimal recognition site is 6 bp, with a likely higher limit of 12 bp. This range of site size yields a *K*_*m•app*_ range from 34.5 to 69.0 nM. The calculated *k*_*cat•app*_ for MMTV intasomes is 0.01 min^-1^. Error bars indicate the standard deviation of at least 3 independent experiments performed with at least 2 independent MMTV intasome preparations.

Because the integration reaction kinetics were linear when the CI products were below 20 fmol we performed a Michealis-Menten substrate concentration analysis (Figure 2B). Sequencing of integration sites in genomic DNA indicated that MMTV has no apparent DNA sequence preference (39). Structural studies have suggested that the MMTV intasome may occupy a 6-12 bp footprint (8). Together these observations suggest an ∼30 fold excess of target sites at these integration assay conditions. Moreover, retroviral intasomes do not turn over. As might be expected, we found that the MMTV intasome substrate-dependent kinetic data fit extremely well to saturable enzymes with a deceleration component at elevated substrate concentration resulting from the lack of enzyme turnover (52) (Figure 2B). We calculated an apparent *k*_*cat•app*_ (0.01 min^-1^) that reflects the bulk enzyme rate of integration catalysis (Figure 2B). Because the occupied site size is not precisely known we could only calculate a range for the *K*_*m•app*_ (34.5 - 69.0 nM). These results echo previous studies that have suggested retroviral integration is inefficient (25,27). Since an infected cell is likely to harbor a single integration complex capable of catalyzing integration, the cellular concentration will be well below nM suggesting that successful target site identification will be limiting (28).

### MMTV intasome aggregation can be reduced by altering solvent components

Purified PFV intasomes appear to readily aggregate in solution resulting in substantial loss of integration activity (29). This aggregation at least partly explains the lack of PFV integration activity after 5 min at 37° C. Single-molecule total internal reflection fluorescence (smTIRF) imaging employs an OSS to reduce photobleaching of fluorophores by reactive oxygen species (ROS) (43,44,53). Protocatechuate 3,4-dioxygenase (PCD) uses protocatechuic acid (PCA) as a substrate to effectively scavenge free oxygen, which contributes to ROS-induced photobleaching. Interestingly, PCA prevented aggregation of PFV intasomes and enhanced integration efficiency (29). Acetylated bovine serum albumen (BSA) also enhanced PFV integration efficiency, presumably by stabilizing the PFV intasome similar to other enzymes in solution (54,55).

We evaluated the effects of PCA and acetylated BSA on MMTV integration and found that neither significantly enhanced MMTV integration when included as a buffer component (Supplemental Figure S3). Moreover, elevated concentrations of PCA appeared to reduce MMTV intasome CI activity, though this difference did not reach statistical significance. We also determined that pre-incubation of MMTV intasomes with acetylated BSA for 30 min at 37° C before the addition of target DNA did not significantly increase integration efficiency, consistent with little or no stabilization of the intasomes (data not shown).

We evaluated the aggregation of MMTV intasomes in reactions that included CaCl_2_ (1 mM) carried over from the purification. This analysis relied on centrifugal precipitation of high molecular weight aggregates. We compared the relative quantity of precipitated aggregates in the absence or presence of additives. A significant reduction in aggregation was observed with the addition of acetylated BSA (47%, p = 0.003; Figure 3A, lane 7 compared to lane 1). This result parallels studies with PFV intasomes and suggests acetylated BSA fundamentally alters MMTV intasome solvation (29,54,55). MMTV intasome aggregation was reduced by 34% with additional CaCl_2_ (10 mM, p = 0.008; Figure 3A). MgSO_4_ also reduced the aggregation of MMTV intasomes by 29% (20 mM, p = 0.05). Elevated NaCl concentrations similarly reduced MMTV intasome aggregation (Figure 3B). However, MMTV intasome integration activity peaked at physiological monovalent salt concentrations (100-150 mM NaCl, Figure 1D) despite the relatively higher aggregation. It is notable that the prevention of MMTV intasome aggregation at the higher NaCl concentrations was similar to that achieved in the presence of 0.2 µg/ml acetylated BSA.

**Figure 3.**
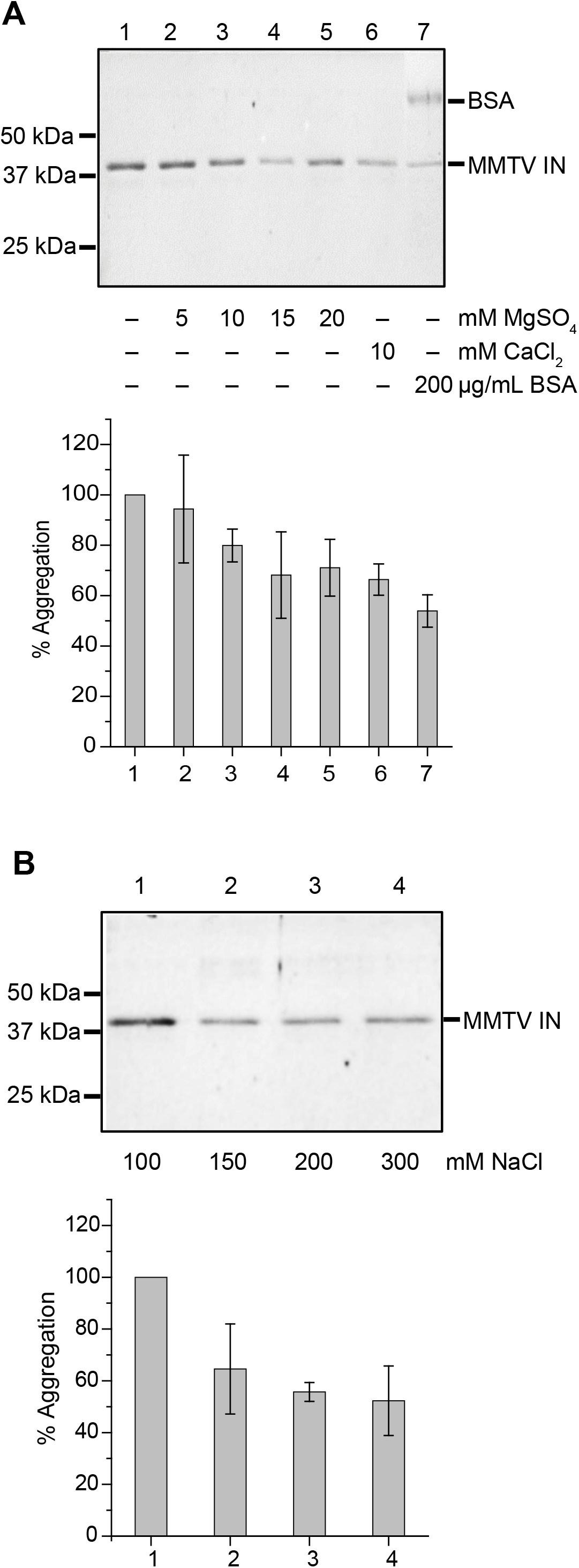
MMTV intasome aggregation. MMTV intasomes were incubated for 1 h at 37°C in integration reaction buffer with indicated concentrations of components. All aggregation reactions include 1 mM CaCl_2_ carried over from the purification buffer. Aggregates were pelleted by centrifugation and analyzed by denaturing PAGE stained with Coomassie blue. (**A**) The percentage of precipitated MMTV IN (% Aggregation) is relative to the control lane (lane 1). Increasing concentrations of MgSO_4_ led to decreased aggregation of MMTV intasomes. Addition of 10 mM CaCl_2_ (final concentration 11 mM CaCl_2_) reduced aggregation, similar to the highest concentrations of MgSO_4_. Addition of acetylated BSA was the most effective at preventing MMTV intasome aggregation. (**B**) MMTV intasome aggregation in the presence of 20 mM MgSO_4_ was assayed with a titration of NaCl. NaCl concentrations >100 mM also led to a decrease of MMTV intasome aggregation. The difference in aggregation between 100 mM and 200 or 300 mM NaCl was significant (p < 0.05), but there was no significant difference between aggregation in 150, 200, or 300 mM NaCl. Error bars indicate the standard deviation of at least 3 independent experiments performed with at least 2 independent MMTV intasome preparations.

### Octameric MMTV intasomes display slower strand transfer kinetics and DNA diffusion than tetrameric PFV intasomes

Retroviral intasomes covalently join each end of the viral cDNA to the host DNA via two independent strand transfer reactions. Single-molecule magnetic tweezers (smMT) analysis was utilized to determine the real-time strand transfer kinetics of MMTV intasomes. In this system a target DNA is tethered at one end to a passivated flow-cell surface by multiple contact points along the backbones of both DNA strands, while the second end is tethered by multiple contact points also along the backbones of both DNA strands to a paramagnetic bead (PMB) (28). Counter-clockwise turns of a neodymium magnet at low magnetic field force (0.3 pN) induced negative supercoils which are a preferred substrate for MMTV intasomes. The time between the two intasome-catalyzed strand transfers (*τ*_ST_) may be distinguished with smMT by tracking the vertical and horizontal motions of individual PMBs (Figure 4A; Supplementary Figure S4A). The first strand transfer is visualized as the PMB position changes on the z-axis when the MMTV intasome has nicked the DNA releasing the negative supercoils. The second strand transfer is observed when the intasome nicks the second DNA strand leading to a double strand break and the PMB position is altered in all axes. MMTV intasomes displayed a *τ*_ST•MMTV_ that was nearly 3 fold longer than the *τ*_ST•PFV_ reported for PFV intasomes (Figure 4B,□ *τ*_ST•MMTV_ = 1.31 ± 0.12 s; *τ*_ST•PFV_ = 0.47 ± 0.05 s; mean ± s.e.) (28). Moreover, the maximum time observed between MMTV intasome strand transfers was 8 s (N = 41), while the maximum time observed for PFV intasome strand transfers was 2.5 s (N = 38). The relaxation time (*τ*_RE_) that measures the release of the supercoils from the target DNA following the first strand transfer was extracted from the integration data (Supplementary Figure S4B, 0.21 ± 0.01 s), which is similar to previous results with the PFV intasome as well as nickase-induced supercoil relaxation (28). These results are consistent with the conclusion that the strand transfer kinetics of MMTV intasomes are intrinsically slower than PFV intasomes, but this difference is not a result of MMTV intasome interference with the fundamental kinetics of DNA supercoil relaxation. These studies are unable to distinguish whether the longer MMTV strand transfer time is a result of the larger octameric intasome, the longer distance between strand transfer sites on the target DNA (6 bp for MMTV versus 4 bp for PFV), or some other unknown catalytic factors.

**Figure 4.**
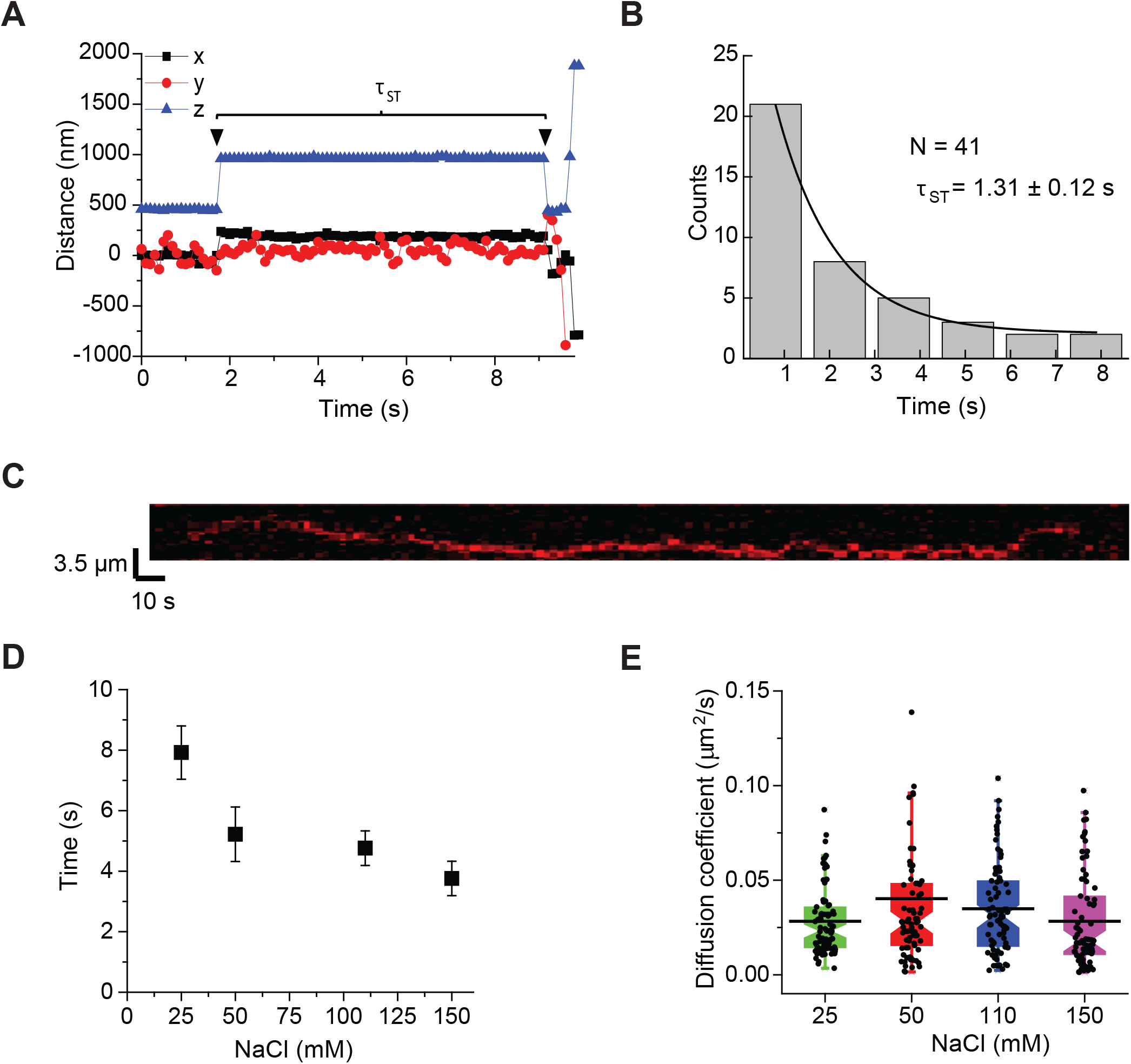
MMTV intasome stand transfer kinetics and search dynamics. (**A**) The time between the two strand transfers (*τ*_ST_) of MMTV vDNA was determined by single-molecule magnetic tweezers. A linear DNA was attached at multiple points to a surface and a paramagnetic bead. Magnets above the microscope stage introduced negative supercoils to the DNA, effectively reducing the apparent height of the bead. The beads were tracked in the x-, y-, and z-axes. The first strand transfer event introduces a nick in the target DNA releasing the supercoils, and changing the z-axis position (blue triangle). The second strand transfer introduces another nick and releases the bead, seen as changes in x-, y-, and z-axes. A representative integration event is shown. The *τ*_ST_ is the time between the first and second strand transfer events. (**B**) The *τ*_ST_ measured for 41 independent integration events was used to construct a histogram. For MMTV intasomes *τ*_ST_ is 1.31 ± 0.12 s. (**C**) Fluorescently labeled MMTV intasomes moving on linear DNA were tracked by TIRF microscopy. A 24 kb linear DNA was attached at both ends to a flow cell surface. 50-100 pM fluorescently labeled MMTV intasomes were added to the flow cell. A representative kymograph of a single MMTV intasome in association with target DNA is shown; the x axis of the trace is time and the y axis is the length of the target DNA. (**D**) The lifetime (*τ*_ON_) of MMTV intasomes in association with target DNA was measured at multiple concentrations of NaCl. Increasing NaCl concentrations had an inverse relationship with the *τ*_ON_ of the intasomes. (**E**) Diffusion coefficients of MMTV intasomes were measured at multiple NaCl concentrations. The diffusion coefficients were similar, indicating a random walk search of ∼2.5 kb at 110 mM NaCl. Error bars indicate the standard deviation of at least 3 independent experiments performed with at least 2 independent MMTV intasome preparations.

Previous studies with PFV intasomes suggested that the rate-limiting step for integration was the DNA search process (28). We used real-time smTIRF microscopy to visualize single MMTV intasomes on a 24 kb *λ*-based target DNA, as previously described (28). Numerous single particles were observed moving along the DNA (Figure 4C; Supplementary Figure S5A; Supplementary Movies 1-4). The lifetime of the MMTV intasome-DNA interaction decreased with increasing salt concentration consistent with increasing ionic shielding of the DNA backbone from protein binding activity (Figure 4D; Supplementary Figure S5B, S6A-D). Within the physiological ionic window, the lifetime of MMTV intasomes on the target DNA appeared to be 4.6 ± 0.6 s, approximately twice as long as PFV intasomes (2.1 ± 0.1 s) (28).

The diffusion coefficient of MMTV intasome particles on the DNA (D_MMTV_ = 0.035 ± 0.02 μm^2^/s, 110 mM NaCl) was approximately 2 fold slower than PFV intasomes (D_PFV_ = 0.082 ± 0.005 μm^2^/s, 110 mM NaCl) (28). These observations are consistent with an increased Stokes’ drag associated with the larger octameric size of MMTV intasomes compared to the tetrameric PFV intasomes. We determined that the diffusion coefficient was constant over a range of ionic strength, consistent with the conclusion that the MMTV intasome maintains continuous contact with the DNA backbone (Figure 4E; Supplementary Figure S5B). Unfortunately, there is currently no structure of the MMTV intasome bound to a target DNA, which is minimally required to calculate a rough landscape diffusion free-energy barrier that might suggest a possible rotation-coupled diffusion mechanism along the DNA (56,57). Nevertheless, the diffusion coefficient indicates that the MMTV intasome may interact with approximately 2.5 kb of target DNA during an average 4.60 ± 0.57 s lifetime in 110 mM NaCl, most likely in a rotation-coupled site-search capacity. We did not observe any concerted integration events during 322 searches of smTIRF target DNA. The lack of efficient integration into a linear target DNA is typical for retroviral intasomes (28).

Catalytically inactive intasomes were assembled with the point mutant MMTV IN(D122N) and Cy5 labeled vDNA. These inactive MMTV intasomes displayed a similar lifetime (4.71 ± 0.33 s) and diffusion coefficient (0.033 ± 0.02 μm^2^/s) to wild type MMTV intasomes (Supplementary Figure S5C,D, S6E). These results are consistent with the conclusion that catalysis of strand transfer is independent of DNA binding and diffusion along the target DNA backbone (28).

### MMTV intasomes form filaments on target DNA

At low MMTV intasome concentrations single particles may be observed interacting with and diffusing along a target DNA by smTIRF. However, when the MMTV intasome concentration is increased, particles appear to aggregate and ultimately form nucleoprotein filaments along the entire length of the 24 kb duplex target DNA (Figure 5A). Moreover, at the initiation of these aggregates there is no visible motion of the intasomes along the DNA. Importantly, we observed little if any integration events that would result in DNA breakage and segregation of the two halves into visible globular aggregates (Figure 5A). We examined the stability of these aggregates by centrifugal precipitation analysis. Intasomes reconstituted with wild type IN or IN(D122N) appeared significantly more prone to precipitation in the presence of linear DNA (Figure 5B). Together these observations suggest that MMTV intasomes may spontaneously form stable aggregates that progress to nucleoprotein filaments when in proximity on target DNA.

**Figure 5.**
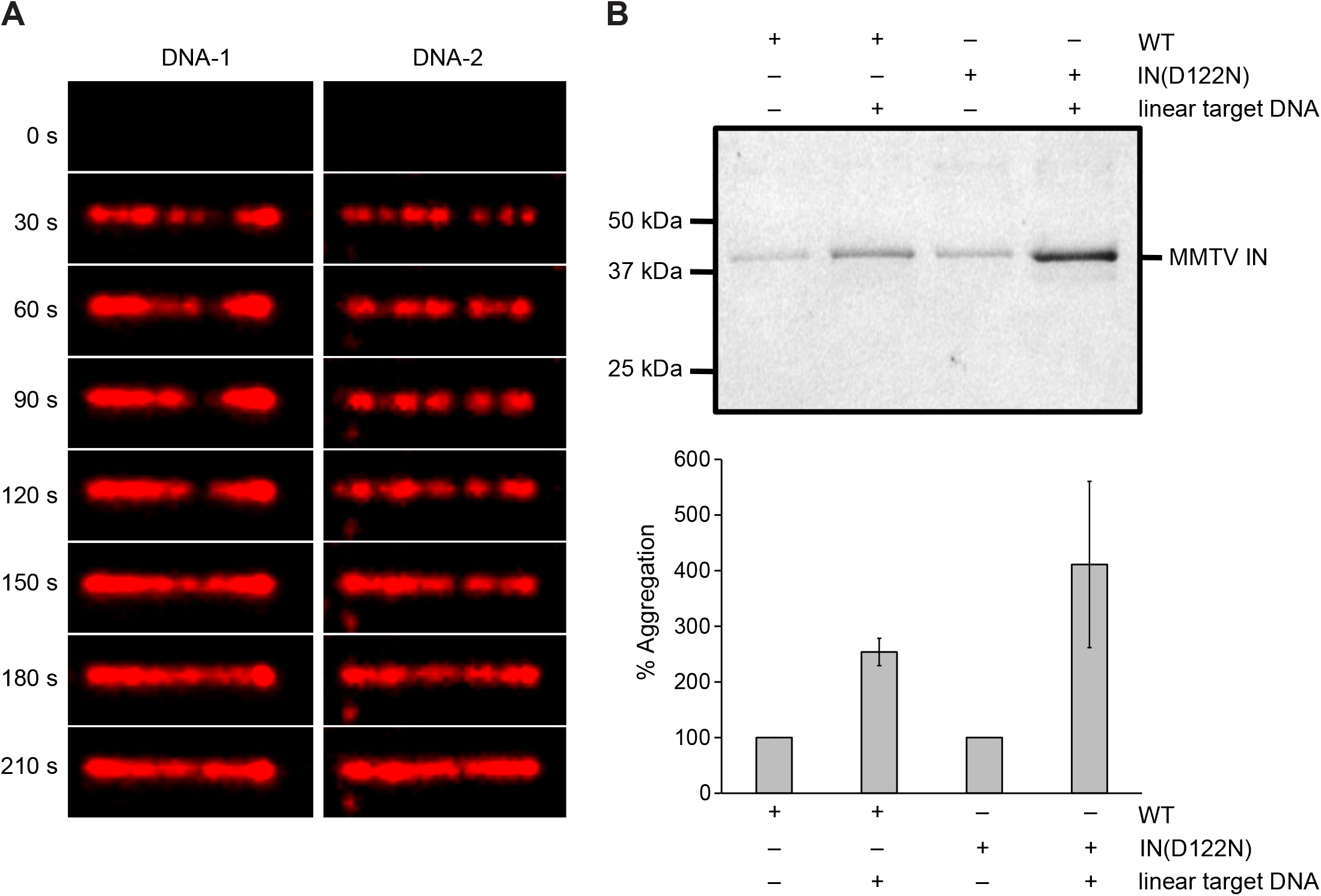
MMTV intasomes are prone to aggregation in the presence of target DNA. (**A**) When 2 nM MMTV intasomes were injected in the flow cell they aggregated on the stretched DNA molecules. The images suggest recruitment and accumulation of additional MMTV intasomes binding the target DNA molecules over time. (**B**) MMTV intasome aggregation was assayed in the absence or presence of 3 kb linear DNA. Wild type and catalytically inactive IN(D122N) intasomes were incubated for 1 h at 37°C in the presence of 110 mM NaCl. Aggregates were pelleted by centrifugation and analyzed by denaturing PAGE. Both wild type and mutant intasomes displayed increased aggregation when linear DNA was present, indicating that aggregation does not require catalysis. The graph represents the mean of three independent experiments performed with at least two independent MMTV intasome purifications.

## DISCUSSION

The tetrameric PFV intasome was the first high resolution retroviral intasome structure described and has served as a model for studies of intasome mediated integration (13,15). Two catalytically active inner PFV IN protomers were shown to position the vDNA ends near a target DNA, while the outer two IN protomers were not completely resolved but appear to play structural roles in stability of the complex (13,15). Several other retroviral intasome structures have been visualized in recent years revealing a spectrum of IN multimers (8,9,11-13,15,17,20,21). A common feature of all these intasome multimers is a CIC (24). The roles of additional IN protomers outside the CIC are largely unknown.

Here we characterized the activity of octameric MMTV intasomes under a variety of biochemical conditions. Importantly, calcium cations are essential for the assembly of stable intasomes because they stabilize the IN-vDNA interaction but are incapable of acting as a cofactor during strand transfer (8,46). These calcium cations must be displaced by magnesium or manganese that can serve as metal ion cofactors capable of coordinating the S_N_2 nucleophilic integration reaction (58). We noted a significant increase in bulk integration products in the presence of the manganese cation (Figure 1). Manganese cations have been shown to alter the catalytic activity of the transposase from bacterial transposon Tn10, while HIV-1 IN displays equivalent activity in magnesium or manganese (46,48,49,59). Similarly, PFV displays equivalent strand transfer activity in both divalent cations, but exhibits increased 3’-end processing in the presence of manganese (47). Since the MMTV intasomes were assembled with oligomers that mimic the 3’-end processed vDNA the increased integration activity with manganese cannot result from enhanced 3’-end processing but rather improved strand transfer. Consistent with this conclusion, manganese also led to increased AI products that can only be formed by strand transfer. Manganese cations display a more relaxed catalytic coordination that may facilitate end-to-end joining of vDNA, non-specific DNA endonuclease activity, and/or non-specific alcoholysis (60-62). However, we regard it more likely that the enhanced catalytic activity exhibited by the manganese cation reflects its ability to displace calcium from the catalytic active site similar to other enzymes (51). These observations highlight the unique behavior of different retroviral IN enzymes to variably employ different divalent cations.

Detailed dynamic processes of PFV intasome association and integration into a target DNA have been visualized by smTIRF and smMT techniques. These previous studies suggested that the PFV intasome binds and moves along a linear target DNA for 2.1 sec by rotation-coupled diffusion (28). Together with an observed diffusion coefficient of 0.08 μm^2^/sec, it was calculated that PFV intasomes interrogate ∼1.6 kb of DNA during every binding event (28). Remarkably, only one of every 100-300 binding events resulted in integration (28). Nevertheless, once an integration site was identified, the time between sequential strand transfer events was 0.47 s on average (28).

Interestingly, the time between MMTV intasome strand transfer events (1.31 s) was ∼3 fold slower than PFV intasomes. The slower strand transfer kinetics may reflect an intrinsically slower strand transfer chemistry or the relatively increased distance between the strand transfer joining points (4 bp for PFV and 6 bp for MMTV), among other possibilities (63). The average lifetime of MMTV intasomes on the DNA (4.6 s) was longer than PFV intasomes. Taken together these observations suggest that MMTV intasomes are catalytically less efficient than PFV intasomes. This conclusion is further mirrored by bulk integration kinetic analysis (Figure 2) that suggested a very slow *k*_*cat•app*_ (0.01 min^-1^) for MMTV integration catalysis and relatively high *K*_*m•app*_ (34.5 - 69.0 nM).

Multiple mechanisms have been described for molecular searches of DNA (64-70). An ionic strength-independent diffusion was observed for both MMTV intasomes and PFV intasomes in agreement with one common search mechanism (28). As might be expected, the diffusion of the octameric MMTV intasome on a target DNA (D_MMTV_ = 0.035 ± 0.02 μm^2^/s) was significantly slower than the tetrameric PFV (D_PFV_ = 0.082 ± 0.005 μm^2^/s). In the absence of a structure with the MMTV intasome bound to target DNA, it is impossible to calculate the rough landscape diffusion constants that would strengthen a rotation-coupled diffusion search model. However, the data is consistent with a model of 1D diffusion in continuous contact with the target DNA (64).

The combination of 1D and 3D mechanisms enhance DNA site search efficiency. Rotation-coupled 1D facilitated diffusion has been observed for the Type II restriction endonuclease EcoRV, which is also able to perform 3D dissociation/reassociation searching (71,72). The single-molecule imaging approaches presented here are unable to determine a role of 3D searching during retroviral intasome interaction with target DNA. Studies of HIV-1 IN predating intasome purification methods suggested a quick commitment to target DNA suggesting that this retroviral IN is incapable of 3D searching (73). However, similar experiments with recombinant PFV IN showed it to be more promiscuous for target DNA, supporting a hypothesis that retroviruses may employ multiple target search mechanisms (47).

There is no known host co-factor for either PFV or MMTV intasomes, unlike lentiviral intasomes that require LEDGF/p75 (8,9,11,74). During infection, PFV and MMTV integration profiles suggest that there is little to no preference for genomic features, such as transcription units or promoters (17,30,37,75,76). This information offers little insight as to how PFV or MMTV intasomes find a target site *in vivo*. Host co-factors, such as LEDGF/p75, that include chromatin and DNA binding domains may dramatically alter the search mechanisms of intasomes. Thus, the influence of a host co-factor search mechanism on intasome search dynamics remains to be explored.

Finally, we have found that increasing the concentration of MMTV intasomes results in the formation of stable aggregates and filaments on the target DNA (Figure 5). We note that HIV-1 pre-integration complexes have been observed to merge and form clusters in the nuclei of primary monocyte derived macrophages (77). Moreover, HIV-1 intasomes that appear to consist of at least a dodecamer of IN subunits also form filaments in solution in the absence of a target DNA (22). This type of aggregation activity suggests that IN associations outside the CIC may reflect both specific and non-specific interactions. It is notable that MMTV intasomes resemble PFV intasomes in DNA search mechanics, strand transfer kinetics, lack of a requisite host co-factor, and ease of recombinant intasome reconstitution. Alternatively, MMTV integration complexes more closely resemble HIV-1 integration complexes in IN multimerization beyond the CIC as well as aggregate/filament formation. These observations appear to suggest that that the MMTV intasome occupies an assembly and catalytic median between the intasomes of PFV and HIV-1. It is likely that comparison of PFV, MMTV, and HIV-1 intasome dynamics will help to parse the mechanical processes that define integration for distinct retrovirus species.

## AVAILABILITY

Video Spot Tracker is available in the GitHub repository (https://github.com/CISMM/video). DiaTrack software is available for download (http://www.diatrack.org/contact.html).

## SUPPLEMENTARY DATA

Supplementary data are available at NAR online.

