## Supplementary Figures for "The mouse mammary tumor virus intasome exhibits distinct dynamics on target DNA"

### Supplementary Figure 1

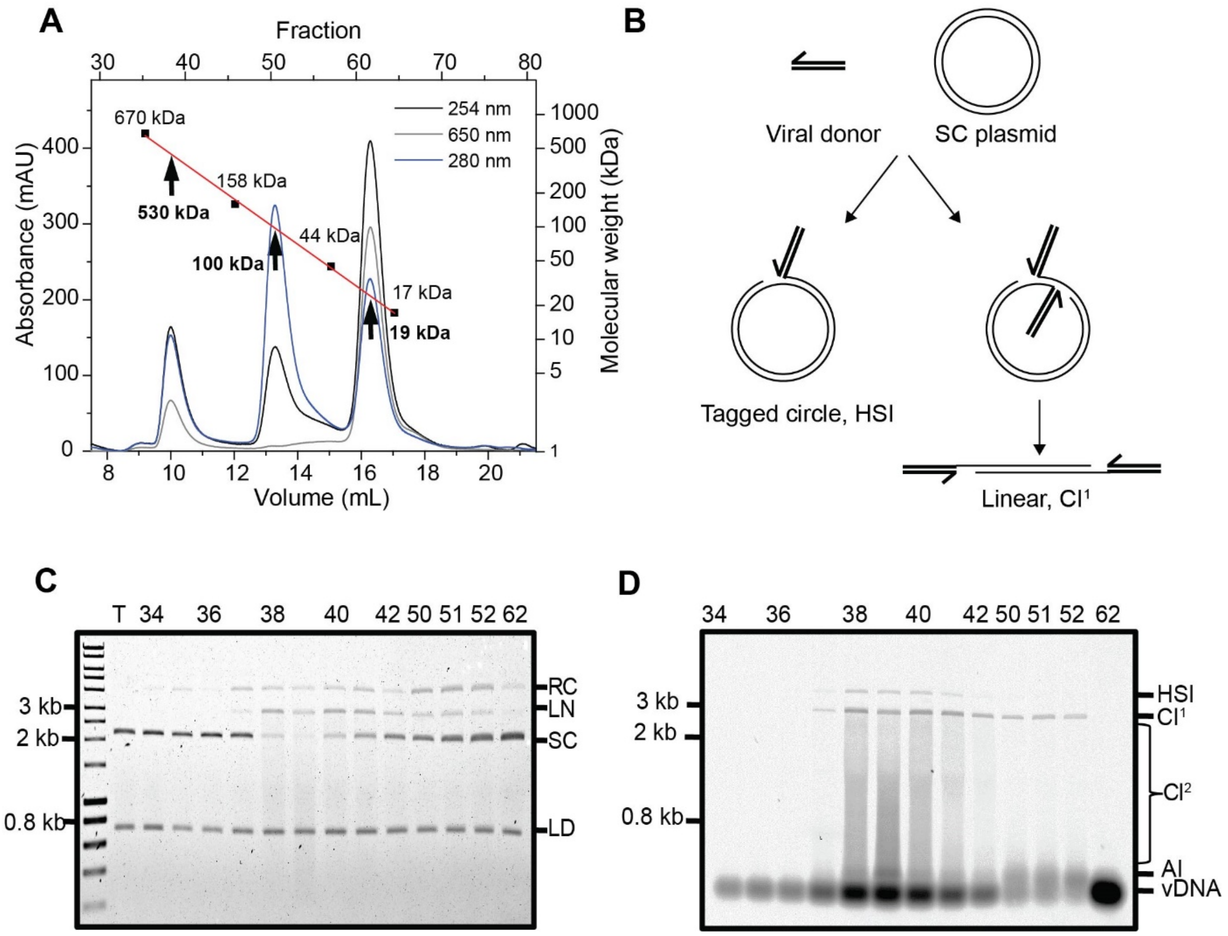


**Supplementary Figure S1. Purification and activity of Cy5 labeled MMTV intasomes.** (**A**) SEC chromatogram of Cy5 labeled octameric MMTV intasome (elution at ~10 mL volume), dimeric MMTV IN (elution at ~13 mL) and Cy5 labeled DNA (elution at ~16 mL). Fraction numbers (top) and elution volume (bottom) are indicated. Fluorophore Cy5 is excited at 650 nm. Protein molecular weight markers (670, 158, 44, and 17 kDa) were profiled in the MMTV intasome buffer and generated a standard curve (red line). The standard curve equation Log10 (MW) = -0.2412 ($V_{elution}$) + 5.1361 ($R^2$ = 0.9712) was used to approximate the molecular weights of the three observed MMTV species (530, 100, and 19 kDa, black arrows). (**B**) Integration of a viral donor DNA (vDNA) converts a supercoiled (SC) plasmid target to several products that may be resolved by agarose gel electrophoresis. Half-site integration (HSI) of a single vDNA to the plasmid results in a tagged circle with the mobility of a relaxed circle (RC). A single concerted integration of both vDNAs to the plasmid yields a product ($CI^1$) with the mobility of linearized plasmid (LN). Additional concerted integration events to $CI^1$ result in a smear of products with mobility between linear plasmid and unreacted vDNA ($CI^2$). (**C**) SEC fractions 34-42, 50-52, and 62 were tested for integration to a 3 kb supercoiled plasmid target. Integration reaction products were separated by agarose gel electrophoresis and imaged by ethidium bromide staining. The ethidium bromide image revealed RC, LN, and SC plasmid as well as a linear DNA loading control (LD). (**D**) The identities of integration products were confirmed by Cy5 imaging of the agarose gel. RC may be the result of HSI or non-specific endonuclease nicking but the presence of Cy5 fluorescence indicates HSI. LN bands correlate with $CI^1$ products. Multiple integration events to a single target DNA generate linear fragments with faster mobility than the linear $CI^1$ product. This smear of $CI^2$ products was more apparent with the Cy5 image. Intermolecular autointegration (AI) uses vDNA as an integration target. The highest integration activity is seen in fractions corresponding to the peak at 10 mL elution volume.

### Supplementary Figure 2

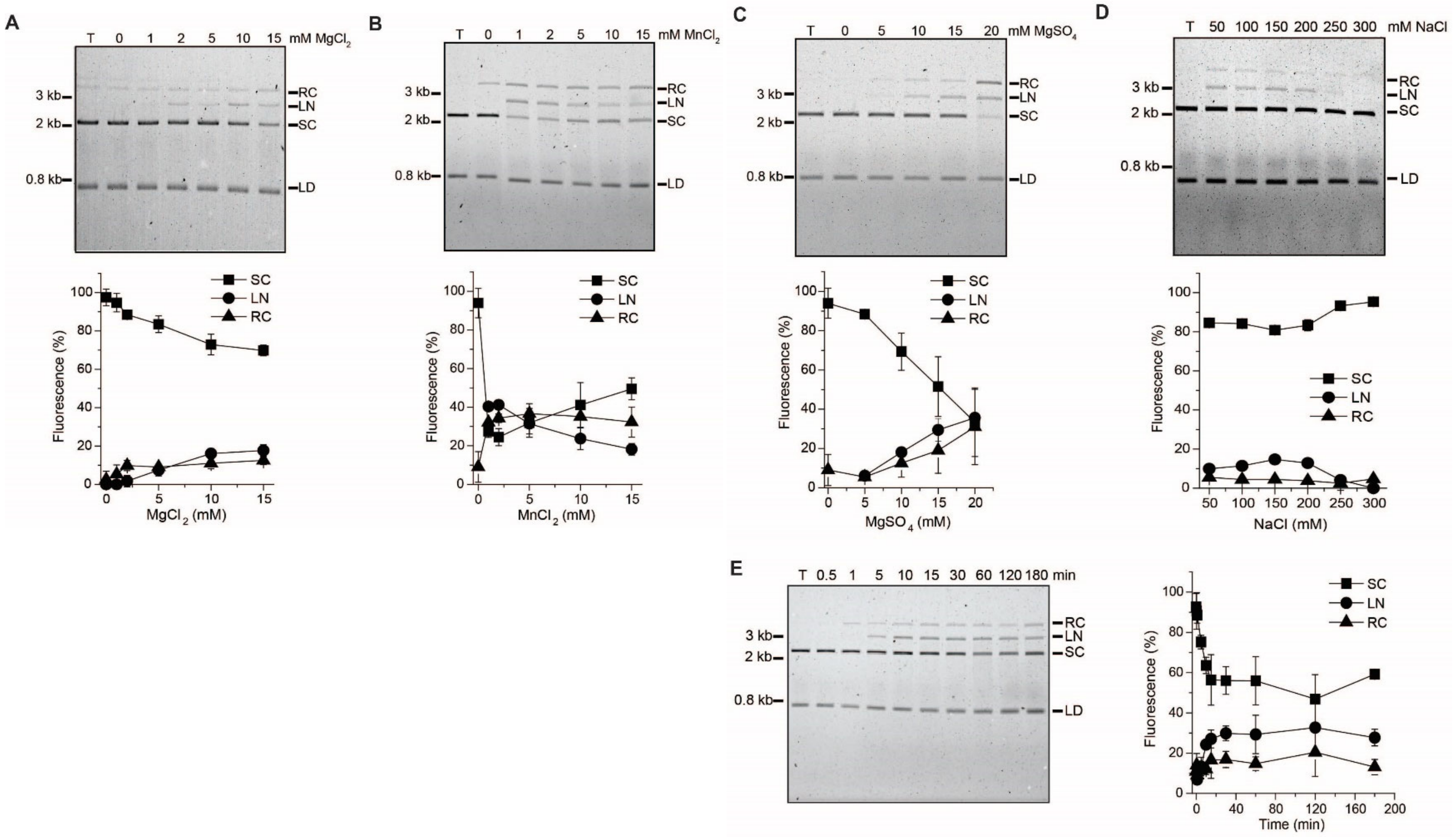


**Supplementary Figure S2. MMTV intasome integration assays with titrations of divalent and monovalent salts and a time course.** Ethidium bromide gel images correspond to the Cy5 images of Figures 1 and 2. Quantitation of the ethidium bromide bands is expressed as the percentage of total fluorescence in the lane. Addition of increasing concentrations of (**A**) $MgCl_2$, (**B**) $MnCl_2$, or (**C**) $MgSO_4$ enhance MMTV intasome integration activity. (**D**) MMTV intasomes displayed maximal activity in the presence of 100-150 mM NaCl; higher concentrations of NaCl reduced MMTV intasome integration efficiency. (**E**) MMTV intasome integration activity increases over time. Graphs represent the mean of three independent experiments performed with at least two independent MMTV intasome purifications. Error bars indicate standard deviation.

### Supplementary Figure 3

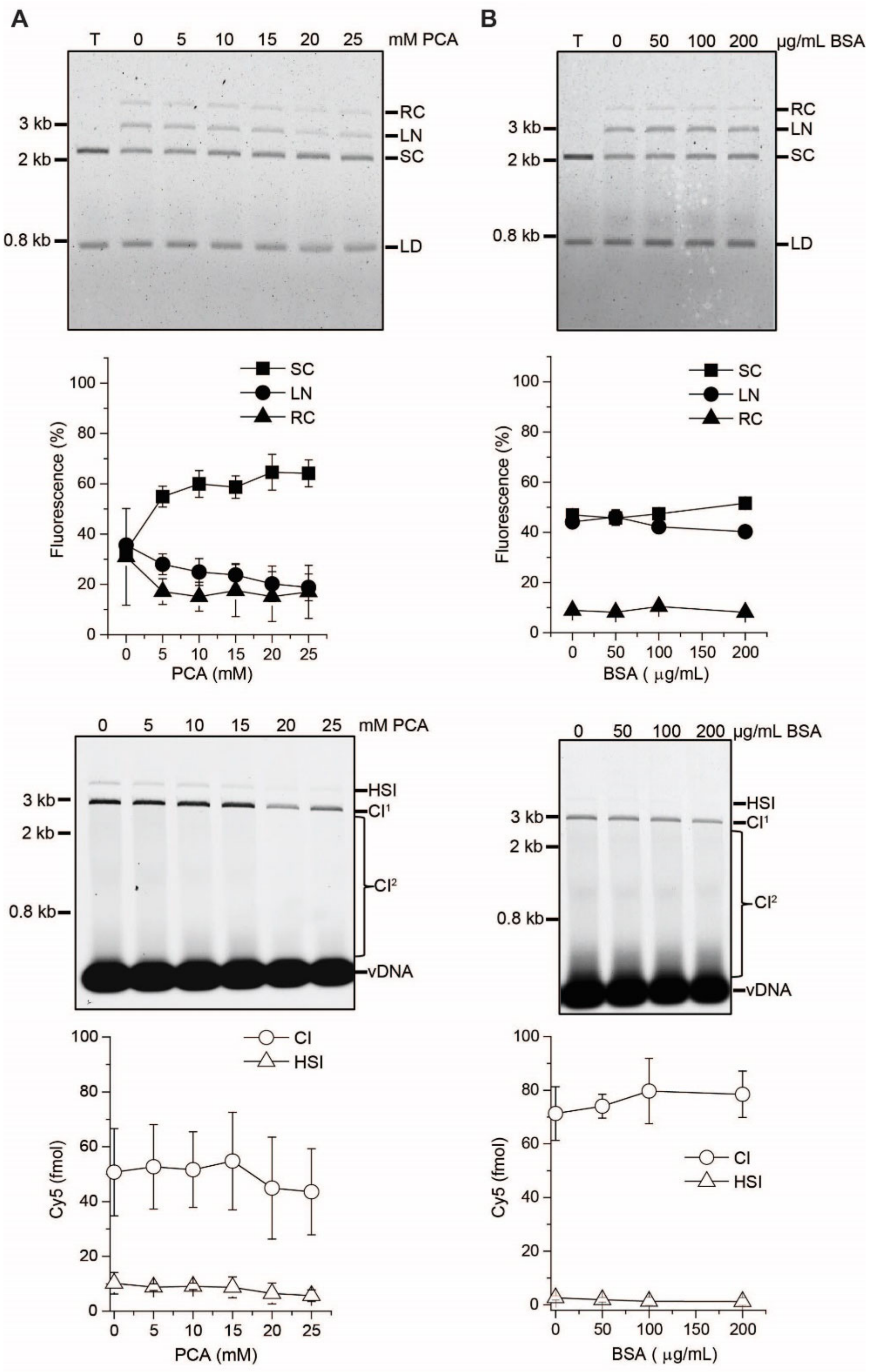


**Supplementary Figure S3. Titrations of PCA or acetylated BSA during MMTV intasome integration assays.** (**A**) Increasing concentrations of PCA reduced MMTV intasome integration activity. Ethidium bromide gel image and quantitation of the ethidium bromide bands is expressed as the percentage of total fluorescence in the lane (top). The percentage of linear (LN) products appeared to decrease while the percentage of unreacted supercoiled (SC) plasmid increased, suggestive of a decrease in concerted integration as PCA concentration increased. The same gel was imaged for Cy5 fluorescence and quantified (bottom). Integration products were quantified by the Cy5 fluorescent signal in each lane (fmol). There were no significant differences in CI products throughout the titration of PCA, suggesting that any effects of this small molecule on MMTV intasomes are subtle. (**B**) Similar to the results with PCA, addition of acetylated BSA showed no apparent effects on MMTV intasome integration efficiency. Graphs represent the mean of three independent experiments performed with at least two independent MMTV intasome purifications. Error bars indicate standard deviation.

### Supplementary Figure 4

**A**

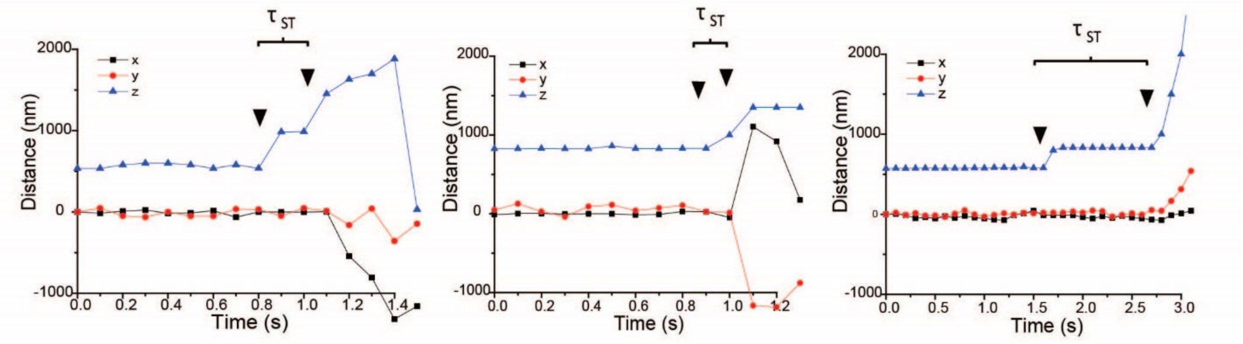


**B**

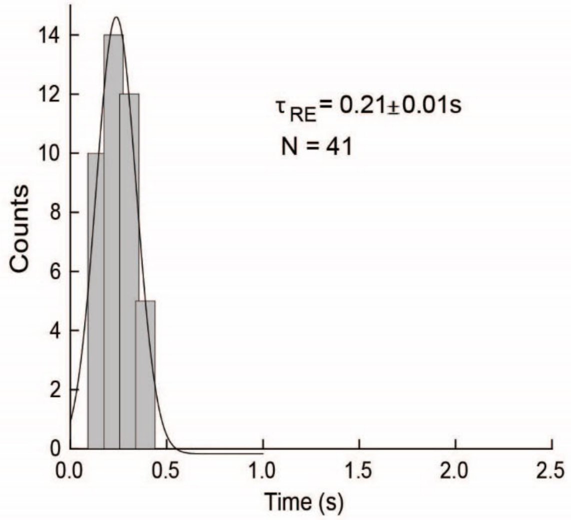


**Supplementary Figure S4. Relaxation time of a supercoiled DNA after the first MMTV strand transfer.** (**A**) A linear DNA was attached at multiple points to a surface and a paramagnetic bead. Magnets above the microscope stage were used to introduce negative supercoils to the DNA, reducing the apparent height of the bead. The first strand transfer nicks the DNA and releases the supercoils, producing a change in the z-axis. The second strand transfer breaks the DNA and the bead leaves the field of view. Three representative integration reactions are shown. (**B**) The relaxation time ($\tau_{RE}$) is the time for the paramagnetic bead to move in the z axis from the supercoiled position to the extended position. The $\tau_{RE}$ is observed immediately after the first strand transfer when a nick releases the supercoils. The $\tau_{RE}$ observed with MMTV intasomes is 0.21 ± 0.01 s (s.e.), consistent with previous $\tau_{RE}$ observations of both PFV intasome and a nicking restriction endonuclease (28).

### Supplementary Figure 5

A

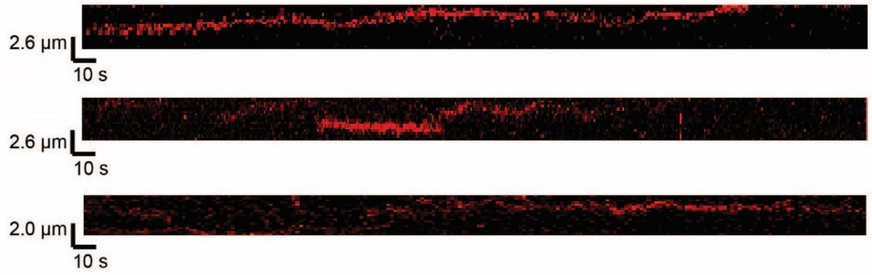


B

| NaCl | $\tau_{ON}$DNA (s) | D (μm²/s) |
|---|---|---|
| 25 mM | 7.92 ± 0.88 | 0.028 ± 0.02 |
| 50 mM | 5.22 ± 0.90 | 0.040 ± 0.05 |
| 110 mM | 4.60 ± 0.57 | 0.035 ± 0.02 |
| 150 mM | 3.76 ± 0.57 | 0.025 ± 0.02 |

C

| | $\tau_{ON}$DNA (s) | D (μm²/s) |
|---|---|---|
| MMTV WT | 4.60 ± 0.57 | 0.035 ± 0.02 |
| MMTV IN(D122N) | 4.71 ± 0.33 | 0.033 ± 0.02 |

D

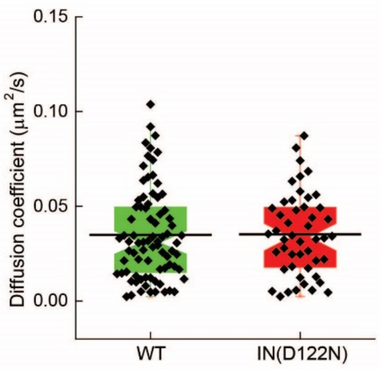


Supplementary Figure S5. MMTV intasome search dynamics with naked DNA. (**A**) Representative traces of a single MMTV intasome in association with target DNA. The x axis of the traces is time and the y axis is the length of the target DNA. (**B**) The average lifetimes ($\tau_{ON}$) of MMTV intasomes in association with DNA and diffusion coefficients at multiple concentrations of NaCl. The $\tau_{ON}$ displays an inverse relationship with NaCl concentration. However, the diffusion coefficients remain constant at multiple NaCl concentrations. (**C**) Catalytically inactive MMTV IN(D122N) intasomes displayed similar $\tau_{ON}$ and diffusion coefficients as wild type intasomes at 110 mM NaCl. (**D**) Diffusion coefficients of wild type and IN(D122N) intasomes in the presence of 110 mM NaCl.

### Supplementary Figure 6

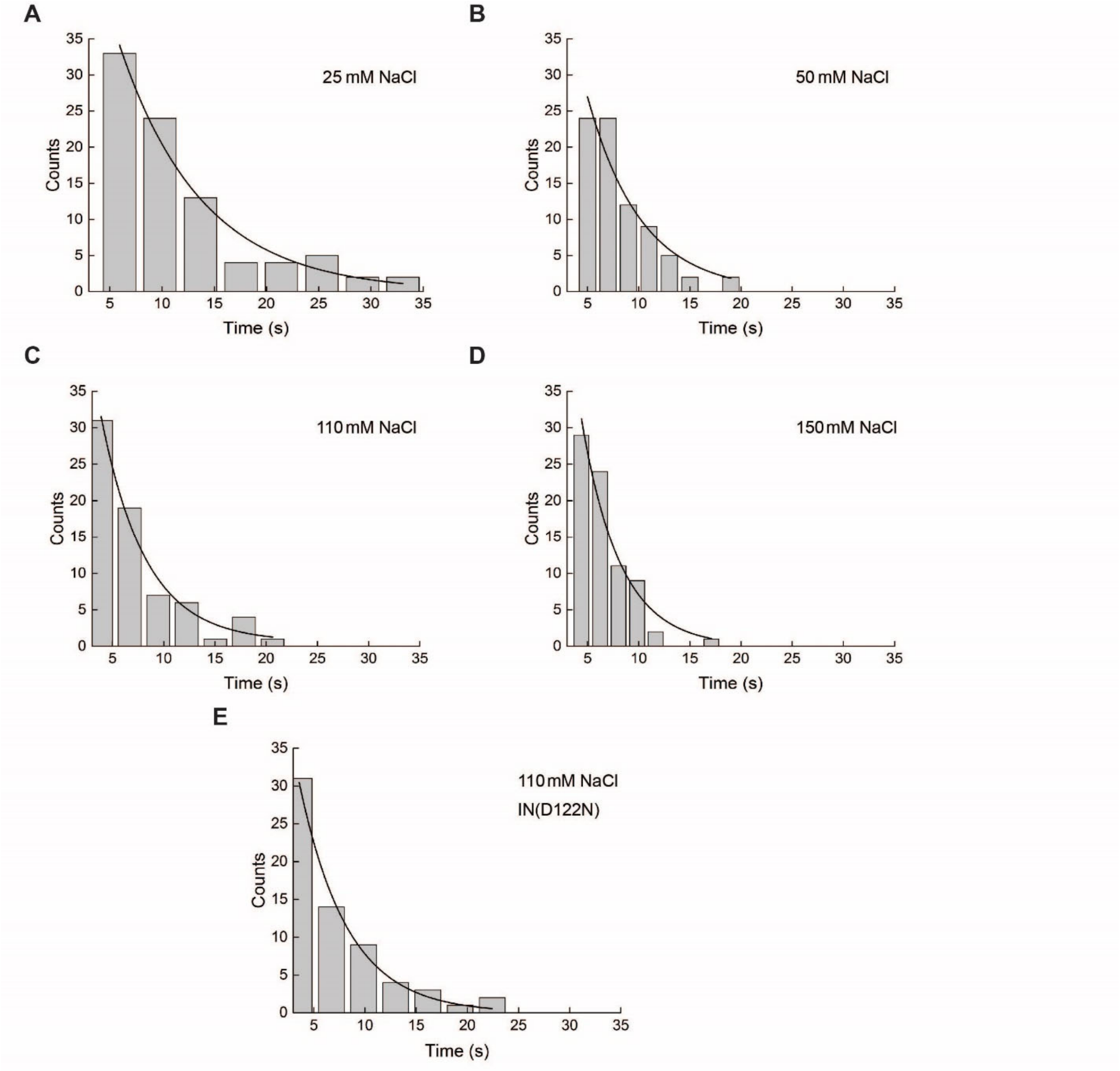


**Supplementary Figure S6. Histograms of observed $\tau_{ON}$ for MMTV intasomes at multiple NaCl concentrations.** The $\tau_{ON}$ for wild type MMTV intasomes is shown for (**A**) 25 mM NaCl, (**B**) 50 mM NaCl, (**C**) 110 mM NaCl, and (**D**) 150 mM NaCl. (**E**) The $\tau_{ON}$ of catalytically inactive IN(D122N) intasomes is shown at 110 mM NaCl. Histogram fits from these plots were used to generate average $\tau_{ON}$ values.

**Supplementary Movies 1-4. MMTV intasomes searching target DNA.** A 24 kb fragment of phage lambda DNA was attached to a surface at both ends. Cy5 labeled MMTV intasomes (green) were imaged at a 250 ms frame rate for 1200 s. Following video capture, the DNA was stained with Sytox and imaged to visualize the 24 kb target DNA (magenta). The Sytox image was overlaid to the MMTV intasome video (co-localization is white). Movies are shown at 3X speed.
